# Calcium Sensing Receptor Common Variants Influence the Effects of Serum Calcium on Coronary Artery Disease Risks

**DOI:** 10.1101/644559

**Authors:** Diane T. Smelser, Fadil M. Hannan, Raghu P.R. Metpally, Sarathbabu Krishnamurthy, David J. Carey, Rajesh V. Thakker, Gerda E. Breitwieser, on behalf of the Regeneron Genetics Center

## Abstract

**Rationale:** The calcium-sensing receptor (CaSR) regulates serum calcium concentrations and common single nucleotide polymorphisms (SNPs) in a carboxyl terminal tri-locus haplotype block contribute to serum calcium variance in the general population. Altered serum calcium concentrations are associated with coronary artery disease (CAD), but direct role for CaSR in CAD remains to be determined.

**Methods:** We evaluated the associations of serum calcium and common *CASR* SNPs or the tri-locus haplotype block with major diseases including CAD in 51,289 patients from the DiscovEHR cohort derived from a single US health care system.

**Results:** Serum calcium concentrations were positively associated with the risk of CAD, and this risk was modified by common *CASR* SNPs. The Ala986Ser SNP was positively associated with hypercalcemia. Carriers of Ala986Ser had a significantly increased CAD risk whereas Arg990Gly carriers had a reduced risk relative to the reference SNP, for those with albumin-corrected serum calcium from 8.5-9.5 mg/dL. In the context of the tri-locus haplotype, the reduced CAD risk conferred by Arg990Gly remained significant. Analysis of the association of common CASR SNPs with CAD risk factors showed Arg990Gly was negatively associated with the CAD risk factor of chronic kidney disease, but independent of alterations in lipids, hemoglobin A1c, or blood pressure.

**Conclusions:** This study compares the common approach of single SNP analysis with the impact of a common variant haplotype block and refocuses attention on the CaSR Arg990Gly SNP which reduces the risk of CAD over a specific range of median albumin-corrected calcium concentrations.

**Précis:** Clinical data and whole exome sequences from a cohort of 51,289 individuals (DiscovEHR) were used to assess the independent contributions of serum Ca^2+^ and *CASR* common variants to cardiovascular diseases including CAD.

## Introduction

The calcium-sensing receptor (CaSR), encoded by the *CASR* gene on chromosome 3q21.1, is a 1078 amino acid family C G protein-coupled receptor (GPCR) that is highly expressed in calcitropic tissues including the parathyroid glands and kidneys, where it regulates serum Ca^2+^ concentrations by influencing parathyroid hormone (PTH) secretion and renal Ca^2+^ excretion.^1^ The CaSR represents a major determinant of extracellular Ca^2+^ homeostasis, and a haplotype block containing a cluster of 3 polymorphisms which localize to the intracellular carboxyl terminus (Ala986Ser, Arg990Gly, and Glu1011Gln) has been shown to contribute to the variance in serum Ca^2+^ in the general population, and to influence urinary Ca^2+^ excretion.^2,3^ The Ala986Ser SNP has been linked to higher serum calcium concentrations in the normal range by <u>G</u>enome-<u>W</u>ide <u>A</u>ssociation <u>S</u>tudies (GWAS).^2^ In contrast, the Arg990Gly SNP has a mild gain-of-function phenotype^4^ linked to lower serum calcium concentrations in the normal range, higher sensitivity to the calcimimetic drug cinacalcet (Sensipar™)^5^, and elevated urinary Ca^2+^ and nephrolithiasis in some, but not all studies.^6^

The CaSR is also expressed in non-calcitropic tissues including heart and arterial vessels^7^, and has been implicated in the pathogenesis of cardiovascular disease (CVD). The common *CASR* SNP Ala986Ser has been associated with an increased prevalence of coronary artery disease (CAD), myocardial infarction (MI) and all-cause cardiovascular mortality in hospitalized patients who had undergone coronary angiography.^8^ However, it is unclear whether the CaSR may influence CVD risk through direct effects on the heart and arterial vessels or by indirectly altering serum Ca^2+^ concentrations. Indeed, serum Ca^2+^ represents a CVD risk factor and calcium supplements, which increase serum Ca^2+^ concentrations, have been independently associated with CVD risk, cardiovascular and all-cause mortality.^9,10^ Moreover, a large-scale Mendelian randomization study found that SNPs in Ca^2+^-regulating genes were significantly associated with both increased serum Ca^2+^ concentrations and CAD; however whilst the *CASR* Ala986Ser SNP had the greatest influence on serum Ca^2+^ concentrations, it was not significantly associated with CAD.^11^ It thus remains to be established whether the cardiovascular effects of serum Ca^2+^ are mediated by, occur independently of or in conjunction with common *CASR* variants. Furthermore, association studies characterizing the influence of *CASR* variants and serum Ca^2+^ on calcitropic and non-calcitropic disorders have been hampered by small samples sizes, and meta-analyses are limited by the use of data from heterogeneous populations. To overcome these difficulties, we utilized clinical, biochemical and whole exome sequences from a single patient cohort comprising 51,289 individuals derived from a stable US population of European origin. We assessed the independent contributions of serum Ca^2+^ and *CASR* common variants to cardiovascular diseases including CAD.

## Methods

### DiscovEHR PATIENT COHORT

The cohort consisted of 51,289 patients with a median age of 61 years (interquartile range 48-72 years) (Table S1), from the Geisinger Health System (GHS) who consented to participate in the MyCode Community Health Initiative^12^, and whose germ-line DNA underwent whole exome sequencing (DiscovEHR).^13^

### CLINICAL DATA

The medians and means of all available serum biochemical and clinical parameters were extracted from the electronic health record (EHR) in a de-identified manner. Patients studied had a median value of 14 years of EHR data (Table S1). We denote serum Ca^2+^ concentrations extracted from the available EHR data for each patient and taken only from outpatient visits as lifetime median (LM_EHR_) serum Ca^2+^. For individuals with available albumin concentrations, median serum Ca^2+^ levels were adjusted, as follows: (adjusted Ca^2+^ (Ca^2+^_alb_) = total serum Ca^2+^ (mg/dL) + 0.8*(4-serum albumin (mg/dL)). Unique International Classification of Disease-9 (ICD9) codes for each de-identified patient were obtained from an approved data broker in accordance with Institutional Review Board approvals.

### EXOME SEQUENCING

Sample preparation, exome sequencing, sequence alignment, variant identification, genotype assignment and quality control steps (allele balance < 0.7, high quality combined allele read depth (AD) of ≥ 8 reads, and per sample genotype quality (GQ) of ≥ 30) have been previously described.^13^

### ASSOCIATION STUDIES

ICD9 codes that were recorded for a minimum of 200 patients (3 or more independent encounters per patient) were grouped into Phenome-Wide Association Study (PheWAS) codes (Phecodes).^14,15^ Non-Europeans were excluded, as was one sample from pairs of closely related individuals up to first cousins. Analyses were adjusted for sex, age, age^2^ and first 4 principal components. Association analyses between CASR SNPs and Phecodes were run with R (3.2) and plotted with GraphPad Prism (V.6).^16^ SAS 9.4 for Windows (SAS Institute, Inc.) was used to analyze associations between common variants or haplotypes and continuous variables, using general linear models (GLM), controlling for age and sex. Pairwise comparisons of haplotypes were performed with least square means, and significant differences between clinical codes, complex phenotypes and/or clinical lab values and genotype were determined by Chi-squared analysis with Yates’ correction, presented as two-tailed p-values. Contributions of *CASR* SNPs or haplotypes to serum Ca^2+^ variations were determined by General Mixed Models, using the albumin-corrected median serum Ca^2+^ levels, corrected for age, sex, and BMI.

## Results

### Association of serum Ca^2+^ concentrations with common *CASR* variants

An exome-wide association study was used to determine the association between mean serum Ca^2+^ concentrations and common *CASR* SNPs in the 51,289 patient DiscovEHR cohort. Results showed significant associations at the *CASR* locus (Figure S1). As previously reported^8^, the strongest positive association was with the Ala986Ser (rs1801725) *CASR* SNP (p=1.6E-70; beta=0.07255), whilst a negative association was identified for Arg990Gly (rs1042636) (p=5.75E-15; beta=-0.0442) (Figure S1). The CaSR carboxyl terminus contains a cluster of SNPs (Ala986Ser, Arg990Gly and Glu1011Gln) in a haplotype block that has been associated with serum Ca^2+^ concentrations.^3,17^ We determined the distribution of haplotypes in the 51,289 DiscovEHR cohort, and assessed their effect(s) on median serum Ca^2+^_alb_. Haplotypes that include the Ser986 allele associate with the highest serum Ca^2+^_alb_ concentrations, whereas haplotypes containing Gly990 alleles associate with the lowest serum Ca^2+^_alb_ concentrations in the normal range (Table 1; P<0.0001). The proportion of serum Ca^2+^ variation accounted for by the Ala986Ser SNP was 6.3%, and by the Arg990Gly SNP was 5.4% (controlling for age, sex, and BMI). None of the haplotypes were significantly associated with serum phosphate or PTH concentrations (Table 1; statistical analysis in Supplemental Table 2).

**Table 1.** Common *CASR* carboxyl terminal haplotype distributions in 51,289 individual DiscovEHR cohort. Patients were sorted by *CASR* haplotype status and presented in order of increasing median serum Ca^2+^_alb_ levels. Sequencing data was not explicitly phased; heterozygous states are indicated by parentheses. Median Ca^2+^ _alb_, PTH and creatinine were averaged over all patients having measurements in the EHR (standard deviation and number of individuals in parentheses). Statistically significant comparisons of median serum Ca^2+^_alb_ are presented in Table S2. There were no statistically significant differences among haplotypes for serum PTH or creatinine.

| Genotype | n | Age (SD) | Male % | Median $\text{Ca}^{2+}_{\text{alb}}$ mg/dL (SD; n) | Serum PTH pg/dL (SD; n) | Serum creatinine mg/dL (SD; n) |
| --- | --- | --- | --- | --- | --- | --- |
| A(R/G)(E/Q) | 301 | 59.2(18.3) | 39.5 | 9.13 (0.27; 105) | 52.0 (29.6; 49) | 0.90 (0.5; 178) |
| ARE/AGE | 5,424 | 58.7(18.1) | 41.3 | 9.16 (0.32; 4,344) | 59.2 (49.1; 1,275) | 0.92 (0.45; 3,135) |
| AGE/AGE | 289 | 58.4(18.3) | 37.0 | 9.17 (0.31; 221) | 69.5 (116.6; 63) | 0.90 (0.61; 145) |
| ARE/ARE | 27,938 | 59.0(17.8) | 40.5 | 9.2 (0.32; 22,855) | 61.9 (75.8; 6,561) | 0.91 (0.43; 16,081) |
| ARE/ARQ | 3,343 | 58.7(17.9) | 41.0 | 9.23 (0.32; 2,752) | 63.63 (55.71; 788) | 0.90 (0.39; 1,884) |
| ARQ/ARQ | 189 | 59.2(20.4) | 38.1 | 9.23 (0.37; 155) | 82.3 (88.7; 54) | 0.93 (0.31; 53) |
| (A/S)(R/G)E | 1,032 | 58.7(17.7) | 40.0 | 9.24 (0.31; 840) | 65.21 (66.57; 230) | 0.88 (0.37; 615) |
| ARE/SRE | 10,910 | 59.2(17.7) | 42.1 | 9.27 (0.33; 8,899) | 62.47 (49.74; 2,577) | 0.91 (0.38; 6,364) |
| (A/S)R(Q/E) | 607 | 58.8(17.8) | 44.8 | 9.28 (0.33; 498) | 71.2 (101.2; 146) | 0.93 (0.50; 329) |
| SRE/SRE | 1,214 | 60.5(16.9) | 39.1 | 9.36 (0.33; 980) | 62.2 (55.9; 311) | 0.94 (0.56; 676) |

**Table 2.** Disease associations for carriers of common *CASR* SNPs. Hetero-+ homozygous individuals were analyzed for each SNP. All ICD9 codes with a minimum of 200 patients, independent of genotype, were assessed. Cases (# cases) were individuals with ≥3 instances of a code, and controls (#controls) were individuals having no instances of the code. Presented are calciotropic and cardiovascular phenotypes with p-value < 0.01, uncorrected for multiple testing as this was an exploratory analysis. Associations with a reduced disease risk are shown in **bold**.

| Code | Code description | O.R.<sup>1</sup> | S.E.<sup>2</sup> | beta<sup>3</sup> | p-value | # cases | # controls |
| --- | --- | --- | --- | --- | --- | --- | --- |
| **A986S (rs1801725)** |  |  |  |  |  |  |  |
| **CALCIOTROPIC PHENOTYPES** |  |  |  |  |  |  |  |
| **275.6** | Hypercalcemia | 1.498 | 0.1005 | 0.4037 | 5.92E-5 | 308 | 28,403 |
| **275** | Disorders of mineral metabolism | 1.275 | .06784 | 0.249 | 0.000342 | 764 | 28,403 |
| **252.1** | Hyperparathyroidism | 1.245 | 0.0854 | 0.22 | 0.01 | 477 | 25,903 |
| **CARDIOVASCULAR PHENOTYPES** |  |  |  |  |  |  |  |
| **443.7** | Peripheral angiopathy | 1.574 | 0.149 | 0.453 | 0.00228 | 138 | 23,387 |
| **250.25** | Type 2 diabetes peripheral circulatory disorder | 1.551 | 0.1543 | 0.439 | 0.0045 | 132 | 17,740 |
| **394** | Chronic rheumatic disease of heart valves | 1.319 | 0.107 | 0.277 | 0.00955 | 304 | 26,320 |
| **427.12** | Paroxysmal ventricular tachycardia | 1.329 | 0.1103 | 0.2845 | 0.0099 | 281 | 20,249 |
| **R990G (rs1042636)** |  |  |  |  |  |  |  |
| **CARDIOVASCULAR PHENOTYPES** |  |  |  |  |  |  |  |
| **411.8** | **Chronic ischemic heart disease** | **0.814** | **0.077** | **-0.218** | **0.0044** | **1,832** | **22,130** |
| **411.1** | **Unstable angina** | **0.564** | **0.203** | **-0.572** | **0.00486** | **298** | **22,130** |
| **411.2** | **Myocardial infarction** | **0.826** | **0.075** | **-0.191** | **0.01** | **1,825** | **22,130** |

1. Odds ratio for code. 2. Standard error of odds ratio. 3. Beta value reflects effect size.

### Association of Common CASR Variants with Major Disease Phenotypes

A phenome-wide association study (PheWAS) using EHR-derived phenotypes^14–16^ identified significant disease associations for patients heterozygous (n=7,691) or homozygous (n=721) for the *CASR* Ala986Ser SNP, including hypercalcemia (p=5.9E-5, beta=0.404) hyperparathyroidism, and disorders of mineral metabolism (Figure 1A, Table 2). Both the Ala986Ser and Arg990Gly SNPs were associated with CVD and the strongest associations for the Ala986Ser SNP include an increased risk of peripheral circulatory disorders and paroxysmal ventricular tachycardia (Figure 1A, Table 2), while the strongest associations for the Arg990Gly SNP showed a reduced risk of chronic ischemic heart disease, unstable angina and myocardial infarction (Figure 1B, Table 2).

**Figure 1.**
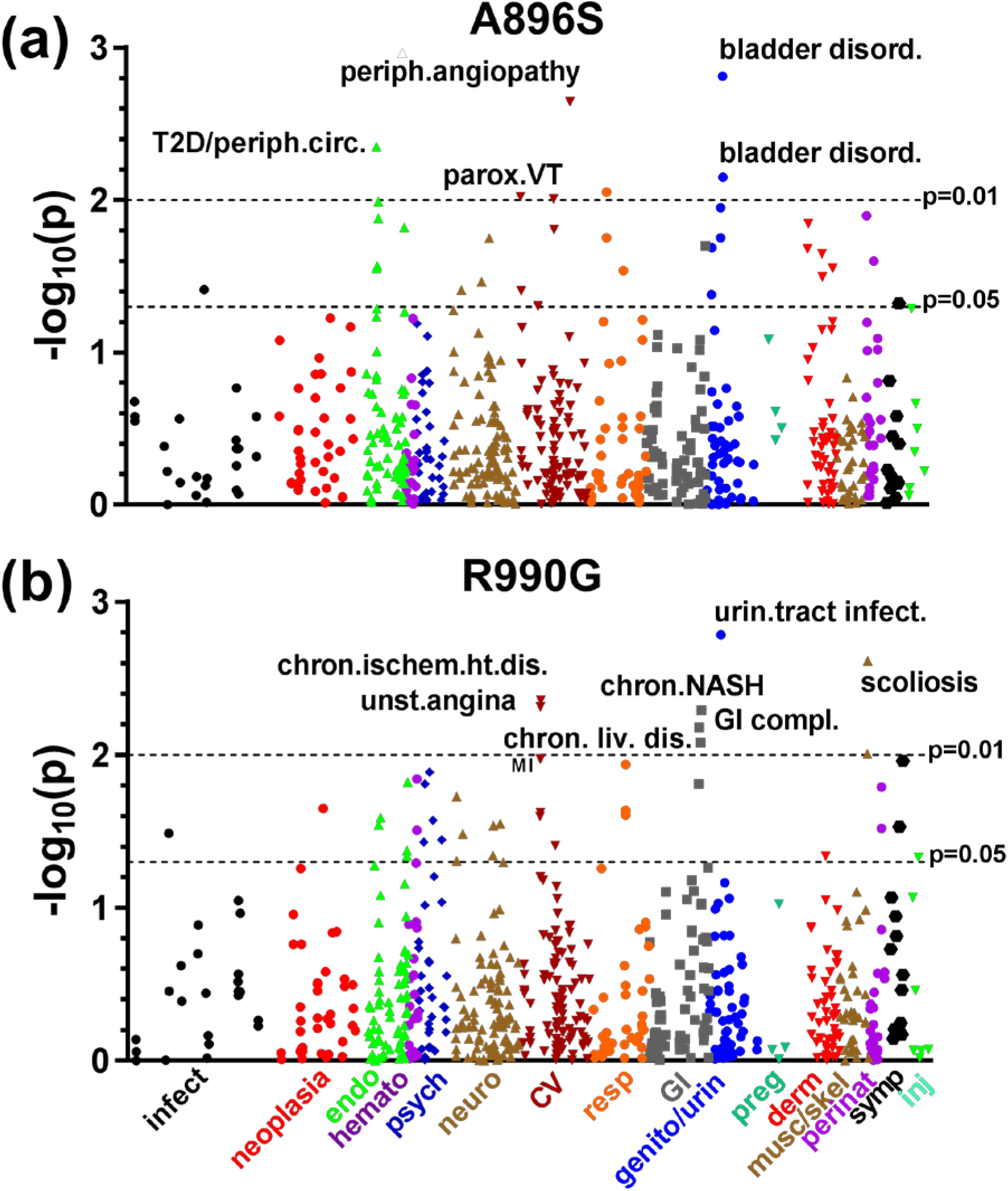
Common *CASR* SNPs are associated with cardiovascular disorders. **A.** PheWAS analysis for those individuals that are homo- or heterozygous for the *CASR* Ala986Ser SNP. Analyzed phenotypes were defined as Phecodes (Methods) with n ≥ 3 calls and at least 200 cases, independent of genotype. All results at p < 0.01 presented in Table 2. **B.** PheWAS results for the Arg990Gly SNP, as described in **A**. All results at p < 0.01 in Table 2.

### Serum Ca^2+^ and *CASR* Variants Modulate Coronary Artery Disease Risk

Associations of the common *CASR* SNPs Ser986 and Gly990 with a range of cardiovascular disease ICD9 codes prompted an in-depth analysis of their respective impacts on CAD, a well-defined clinical phenotype. CAD patients were defined by an EHR history of coronary revascularization, or history of acute coronary syndrome, ischemic heart disease or exertional angina (ICD9 codes 410*-414*) with angiographic evidence of obstructive CAD (>50% stenosis in at least one major epicardial vessel as determined by catheterization).^13,18^ Controls were individuals without any case criteria or any single encounter or problem list diagnosis code indicating CAD.

We first used LM_EHR_ serum Ca^2+^_alb_ ranges as a categorical variable to determine the relative CAD risks of the *CASR* SNPs Ser986 and Gly990 relative to individuals having all reference alleles (WT=Ala986/Arg990/Glu1011). CAD cases were sorted by LM_EHR_ serum Ca^2+^_alb_ range and genotype, and plotted over the range of LM_EHR_ serum Ca^2+^_alb_ from 8.5 to 10.0 mg/dL (Figure 2A). Individuals having one or two Ser986 alleles had a CAD risk in each Ca^2+^ range comparable to the reference allele (WT), while those having one or two Gly990 alleles had a significantly reduced CAD risk relative to Ser986 (O.R. = 0.762, 95% C.I. 0.603-0.96; p=0.021) or WT (O.R. = 0.81, C.I. 0.66-0.98; p=0.034). CAD risks increased with LM_EHR_ serum Ca^2+^_alb_ range for Gly990 (8.5-9.0 mg/dL vs 9.5-10 mg/dL, O.R. = 1.53, C.I. 1.14-2.06; p=0.0053) and WT (8.5-9.0 mg/dL vs 9.5-10 mg/dL, O.R. = 1.3, C.I. 1.13-1.5, p=0.0003). When analyses were corrected for the well-established CAD risk factors of age, sex, and BMI, the Gly990 genotype was shown to be an independent factor which significantly decreased the risk of CAD when LM_EHR_ serum Ca^2+^_alb_ is in the 8.5-9.0 mg/dL range (Figure 2B).

**Figure 2.**
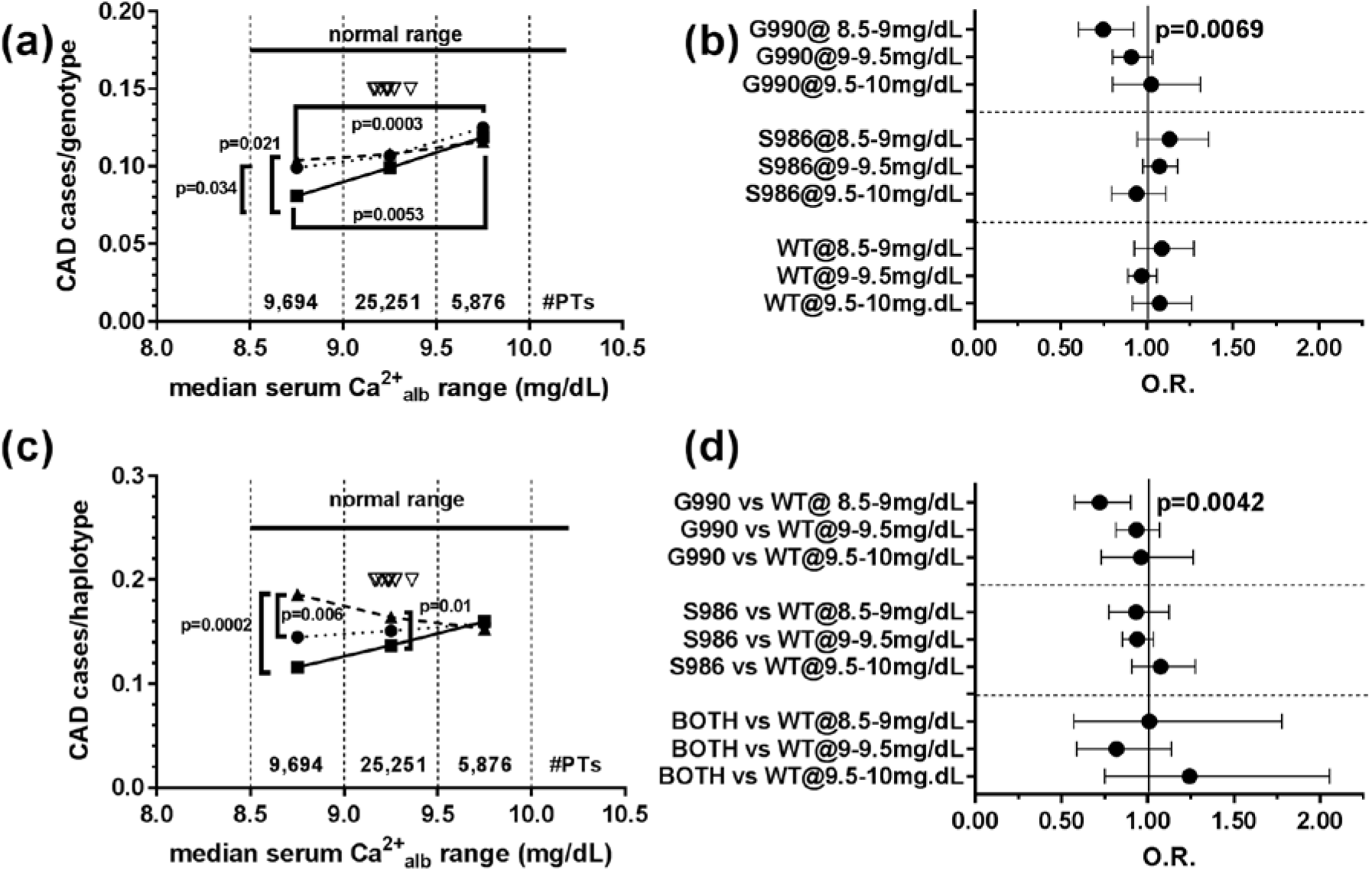
CAD risk as a function of *CASR* SNPs and LM_EHR_ serum Ca^2+^alb. **A.** CAD risk (#CAD patients/total patients of each genotype) was plotted for each Ala986Ser and Arg9990Gly SNPs (WT = Ala986, Arg990, Glu1011) in 0.5 mg/dL ranges of median serum Ca^2+^_alb_ from 8.5-10 mg/dL. The total number of patients in each median Ca^2+^_alb_ range is indicated on the graph. Inverted arrowheads mark the haplotype median Ca^2+^_alb_ values from Table 1; black circles, individuals with reference genotype at all positions; triangles, S986 homo-plus heterozygous individuals; and squares, G990 homo- and heterozygous individuals. Statistical significance was determined by Chi-squared analysis with Yate’s correction using GraphPad Prism (V6). **B.** Analysis of CAD risk as described in **A** for each CASR haplotype (WT haplotype = Ala986, Arg990, Glu1011). For the Ser986- or Gly990-containing haplotypes, other SNPs of the haplotype were reference. Black circles, individuals with reference genotype at all positions, triangles, S986 homo-plus heterozygous individuals, squares, G990 homo- and heterozygous individuals, with all other positions at reference allele. Statistical significance was determined by Chi-squared analysis with Yate’s correction using GraphPad Prism (V6). **C.** Odds ratio (O.R.) analysis of CAD risk as a function of *CASR* SNP and LM_EHR_ serum Ca^2+^_alb_, compared to WT. Plotted are the odds ratios and 95% confidence intervals. Significant odds ratios are indicated with p-values determined by Chi-squared analysis. Analysis was corrected for age, sex, and BMI. Full analyses of the CAD risks and LM_EHR_ Ca^2+^_alb_ dependencies are presented in Supplemental Table 3. **D.** Analysis of CAD risk as in **C.**, using haplotypes, compared with the WT haplotype. Statistical significance (Chi-squared analysis) indicated with p-values. CAD risks as a function of age, sex, and BMI are in Supplemental Table 4.

We next determined whether CAD risks are modified in the context of the carboxyl terminal haplotype (Figure 2C). In this case, one or two Ser986 alleles for patients in the 8.5-9.0 mg/dL LM_EHR_ Ca^2+^_alb_ range significantly increased CAD risk compared with individuals with one or two Gly990 alleles (O.R.= 1.74, C.I. 1.3-2.32, p=0.0002). Likewise, CAD risk was significantly higher for individuals with Ser986 allele(s) than those with the reference haplotype (O.R. = 1.35, C.I. 1.09-1.65, p=0.0058) over the same LM_EHR_ Ca^2+^_alb_ range (Figure 2C). CAD risk was also higher for the Ser986 relative to the Gly990 allele in the serum Ca^2+^_alb_ range from 9.0-9.5 mg/dL (O.R. = 1.23, C.I. 1.05-1.45, p=0.0103) (Figure 2C). When compared with the WT haplotype, those having the Gly990 allele had a reduced CAD risk in the 8.5-9.0 mg/dL serum Ca^2+^_alb_ range (O.R. = 0.805, C.I. 0.66-0.98, p=0.034). CAD risk for patients having median serum Ca^2+^_alb_ levels above 9.5 mg/dL was independent of genotype (Figure 2C). We corrected the analysis of haplotype associated CAD risks for age, sex and BMI (Figure 2D), and results again demonstrated the importance of the Gly990 allele as an independent determinant of reduced CAD risk. Finally, we analyzed the effect of *CASR* haplotype on CAD risk, using LM_EHR_ Ca^2+^_alb_ as a continuous variable. CAD risk is associated with serum Ca^2+^ even for individuals having the reference haplotype, increasing 2.7% per 0.1 mg/dL of serum Ca^2+^, while CAD risk increased by 1.9% per 0.1 mg/dL of Ca^2+^ for carriers of Ser986 alleles, and by 5.5% for each 0.1 mg/dL of Ca^2+^ for carriers of Gly990 alleles. Overall, results demonstrate that presence of the Gly990 allele significantly reduces CAD risk in the 8.5-9.0 mg/dL serum Ca^2+^_alb_ range, and that both serum Ca^2+^ and *CASR* haplotype are strongly associated with CAD risk.

CaSR is highly expressed in the pancreatic islets and kidneys.^19–21^ We therefore determined whether the Ser986 or Gly990 *CASR* SNPs influenced CAD risk through differential effects on the prevalence of type 2 diabetes (T2D) or chronic kidney disease (CKD). Individuals homozygous for the Ser986 SNP had a small but significant increase in the prevalence of T2D in the 51,289 individual DiscovEHR cohort (Figure S2D), but the *CASR* SNPs had no differential effects on other cardio-metabolic risk factors, including dyslipidemia ICD9 calls or blood pressure (Figures S2ABC), glycemia (hemoglobin A1c concentrations, Figure S2E), estimated glomerular filtration rate (Figure S2F) or serum lipids (Figure S3). It should be noted that drug treatments for diabetes and/or dyslipidemia may obscure the full effect of the CaSR variants on these measures. Individuals homozygous for the Gly990 SNP in the background of the reference alleles at both of the other common SNPs in the haplotype had a significantly reduced prevalence of CKD (stages 4 (ICD9 code 585.4) + 5 (585.5) + unspecified (585.9), but excluding end stage renal disease (585.6), Figure 3A), which was reflected in significantly lower serum creatinine levels (Figure 3B). Overall, results suggest that *CASR* common variants modify the effect of serum Ca^2+^ on CAD, either directly and/or by effects in the kidney.

**Figure 3.**
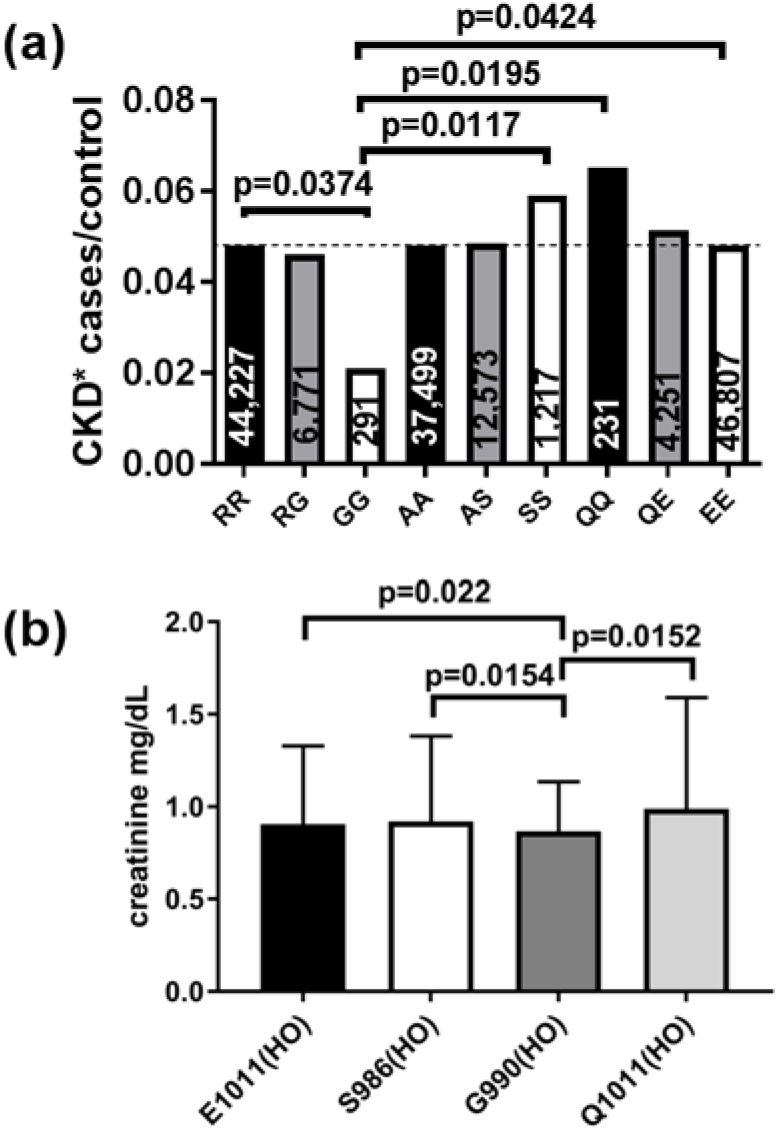
CKD as a function of *CASR* SNPs. **A.** Cases of chronic kidney disease (CKD*, sum of individuals with at least 3 calls of ICD9 codes of 585.5, 585.6, or 585.9, with each patient represented only once at most severe code) as a function of common *CASR* variant genotype. Significance determined by chi-squared analysis (Fisher’s exact test or Chi-squared with Yate’s correction depending on the number of patients being compared). Total number of patients with each genotype are indicated in the bars. Individuals homozygous for the Gly990 SNP had a significantly reduced CKD* risk compared with other genotypes. **B.** Mean of median creatinine values for all individuals homozygous for CASR common haplotypes, irrespective of CAD or CKD status. E1011(HO) denotes the reference haplotype (35,343 individuals), and there were 1059 individuals homozygous for S986, 250 individuals homozygous for G990, and 165 individuals homozygous for Q1011. Significance was determined by two-tailed t-test assuming unequal variance.

## Discussion

The availability of whole exome sequencing coupled with phenotype-rich electronic health record data has provided a unique opportunity to define cardiovascular and other disease associations for common *CASR* variants, and dissect the contribution of serum Ca^2+^ to CAD risk. Our studies have revealed that serum Ca^2+^ concentrations are positively associated with the risk of CAD in patients who are not carriers of CaSR common variants, demonstrating that serum Ca^2+^ likely influences CVD independent of its effects on the CaSR. These findings are consistent with previous observational studies, which have shown that an ~0.1 mmol/L increase in serum Ca^2+^ concentrations is associated with an ∼10% increase in the relative risk of cardiovascular events.^22^ Furthermore, we have dissected the associations of common *CASR* variants with CAD over defined ranges of serum Ca^2+^ concentrations, and demonstrate that common *CASR* variants modify the effect of serum Ca^2+^ on CAD. Our stepwise analysis argues that analysis of single SNPs in a linked haplotype block containing multiple common variants may not take into account all determinants of CAD risk. Therefore, the CASR tri-locus haplotype (Ala986Ser/Arg990Gly/Glu1011Gln) should be used to assess disease risks, as the interaction(s) of risk and protective alleles both need to be taken into account. When considered alone, the *CASR* carboxyl terminal Ser986 SNP significantly increased CAD risk, confirming a previous study^8^, whilst the Gly990 SNP significantly reduced this risk. These opposing effects of the CaSR SNPs on CAD risk occurred at serum Ca^2+^ concentrations in the range between 8.5-9.5 mg/dL, which accounts for the majority (83%) of patients in the CAD analysis. When considered within the tri-locus haplotype, the reduced risk conferred by the Arg990Gly SNP remained significant. Of note, the frequency of the Arg990Gly SNP varies among ethnicities (1000Genomes), and may contribute to observed differential risks of CAD in different populations. The finding that these associations of the *CASR* SNPs with CAD occurred independently of cardio-metabolic risk factors such as blood pressure, serum lipids or hemoglobin A1c concentrations, highlights a potential role for CaSR expressed in the cardiovascular system in the pathogenesis of CAD. Indeed, the CaSR is expressed in the aorta and coronary arteries^23^, and abnormal function of the arterial CaSR has been implicated the pathogenesis of vascular calcification.^24^ Moreover, we have shown that the homozygous Gly990 CaSR SNP is associated with a significant reduction in CKD cases. The relevance of this finding to the pathophysiology of CKD is uncertain, however, it is notable that studies involving rats have shown CaSR to protect against glomerular disease.^25^ Moreover, as CKD represents a major cardiovascular risk factor, the reduction in CKD may in part explain why the Gly990 CaSR SNP is associated with a reduced risk of CAD. Furthermore, the Ser986 CaSR SNP is associated with an increase in cases of type 2 diabetes, and these findings are in keeping with a reported study involving renal transplant recipients, which showed the Ser986 SNP to be associated with increased serum glucose concentrations.^26^

The approaches used in this study have notable benefits and a few limitations. Combining WES and quantitative clinical measures from a single large healthcare system allowed unequivocal definitions of cases and controls, improving the power to discern associations. Another benefit is the ability to study a broad range of clinical conditions in an unbiased manner. The de-identified nature of the data confers some limitations, including a lack of specialized clinical data and/or testing to support particular diagnoses, compounded by the inability to call back patients for additional phenotyping. These limitations promoted a focus on the CAD phenotype, which was validated by angiographic evidence of obstructive CAD^13,18^. Finally, we did not explicitly address whether patients were on supplemental calcium ± vitamin D, which may influence serum Ca^2+^ concentrations, but rather focused on outpatient EHR-based median Ca^2+^_alb_ levels to assess CAD risks. The large numbers of patients and extensive clinical measures did, however, allowed for statistically significant findings, and the present results argue for the importance of including *CASR* haplotypes in re-analyses of renal failure studies that use cinacalcet. Finally, large independent cohorts combining WES and granular EHR data are currently unavailable for replication of the observed associations. As other population sequencing efforts such as the UK Biobank and the National Institutes of Health Precision Health Initiative (All of Us) come to fruition, we expect replication to be possible. The present study can also be followed-up with an in-depth prospective study with specifically recruited patients who can undergo more rigorous testing.

In conclusion, common *CASR* SNPs within the carboxyl terminal tri-locus haplotype are significantly associated serum Ca^2+^ and with cardiovascular diseases, and modify the effects of serum calcium on CAD risk, independently of other risk factors.

## Acknowledgements

We thank the Phenotype Core at Geisinger for providing de-identified clinical data, and the patients who contributed to MyCode to make these studies possible. We thank Dr. Janet Robishaw for comments on an early version of the manuscript. This work is supported by a Wellcome Trust Investigator Award, National Institute for Health Research Senior Investigator Award, and National Institute for Health Research Oxford Biomedical Research Centre Programme (Prof. Thakker).

## References

1. Hannan FM, Babinsky VN, Thakker RV. Disorders of the calcium-sensing receptor and partner proteins: insights into the molecular basis of calcium homeostasis. J Mol Endocrinol. 2016;57:R127–R142.

2. Kapur K, Johnson T, Beckmann ND, Sehmi J, Tanaka T, Kutalik Z, Styrkarsdottir U, Zhang W, Marek D, Gudbjartsson DF, Milaneschi Y, Holm H, Diiorio A, Waterworth D, Li Y, Singleton AB, Bjornsdottir US, Sigurdsson G, Hernandez DG, Desilva R, Elliott P, Eyjolfsson GI, Guralnik JM, Scott J, Thorsteinsdottir U, Bandinelli S, Chambers J, Stefansson K, Waeber G, Ferrucci L, Kooner JS, Mooser V, Vollenweider P, Beckmann JS, Bochud M, Bergmann S. Genome-wide meta-analysis for serum calcium identifies significantly associated SNPs near the calcium-sensing receptor (CASR) gene. PLoS Genet. 2010;6:e1001035.

3. Scillitani A., Guarnieri V, De Geronimo S, Muscarella LA, Battista C, D’Agruma L, Bertoldo F, Florio C, Minisola S, Hendy GN, Cole DE. Blood ionized calcium is associated with clustered polymorphisms in the carboxyl-terminal tail of the calcium-sensing receptor. J Clin Endocrinol Metab. 2004;89:5634–5638.

4. Vezzoli G, Terranegra A, Arcidiacono T, Biasion R., Coviello D., Syren ML, Paloschi V, Giannini S., Mignogna G, Rubinacci A, Ferraretto A, Cusi D, Bianchi G, Soldati L. R990G polymorphism of calcium-sensing receptor does produce a gain-of-function and predispose to primary hypercalciuria. Kidney Int. 2007;71:1155–1162.

5. Jeong S, Kim IW, Oh KH, Han N, Joo KW, Kim HJ, Oh JM. Pharmacogenetic analysis of cinacalcet response in secondary hyperparathyroidism patients. Drug Design, Development and Therapy. 2016;10:2211–2225.

6. Egbuna Ol, Brown EM. Hypercalcaemic and hypocalcaemic conditions due to calcium-sensing receptor mutations. Best Pract Res Clin Rheumatol. 2008;22:129–148.

7. Toka HR, Pollak MR. The role of the calcium-sensing receptor in disorders of abnormal calcium handling and cardiovascular disease. Curr Opin Nephrol Hypertension. 2014;23:494–501.

8. März W, Seelhorst U, Wellnitz B, Wellnitz B., Tiran B, Obermayer-Pietsch B, Renner W, Boehm BO, Ritz E, Hoffmann MM. Alanine to serine polymorphism at position 986 of the calcium-sensing receptor associated with coronary heart disease, myocardial infarction, all-cause, and cardiovascular mortality. J Clin Endocrinol Metab. 2007;92:2363–2369.

9. Thiele I, Linseisen J, Meisinger C, Schwab S, Huth C, Peters A, Perz S, Meitinger T, Krinenberg F, Lamina C, Thiery J, Koenig W, Rathmann W, Kaab S, Then SC, Seissler J, Thorand B. Associations between calcium and vitamin D supplement use as well as their serum concentrations and subclinical cardiovascular disease phenotypes. Atherosclerosis. 2015;241:743–751.

10. Challoumas D, Cobbold C, Dimitrakakis G. Effects of calcium intake on the cardiovascular system in postmenopausal women. Atherosclerosis. 2013;231:1–7.

11. Larsson SC, Burgess S, Michaelsson K. Association of genetic variants related to serum calcium levels with coronary artery disease and myocardial infarction. JAMA. 2017;318:371–380.

12. Carey DJ, Fetterolf SN, Davis FD, Faucett WA, Kirchner HL, Mirshahi U, Murray MF, Smelser DT, Gerhard GS, Ledbetter DH. The Geisinger MyCode community health initiate: an electronic health record-linked biobank for precision medicine research. Genet Med. 2016;18:906–913.

13. Dewey FE, Gusarova V, O’Dushlaine C, Gottesman O, Trejos J, Hunt C, Van Hout CV, Habegger L, Buckler D, Lai KM, Leader JB, Murray MF, Ritchie MD, Kirchner HL, Ledbetter DH, Penn J, Lopez A, Borecki IB, Overton JD, Reid JG, Carey DJ, Murphy AJ, Yancopoulos GD, Baras A, Gromada J, Shuldiner AR. Inactivating variants in ANGPTL4 and risk of coronary artery disease. N Engl J Med. 2016;374:1123–1133.

14. Verma A, Leader JB, Verma SS, Frase A, Wallace J, Dudek S, Lavage DR, Van Hout CV, Dewey FE, Penn J, Lopez A, Overton JD, Carey DJ, Ledbetter DH, Kirchner HL, Ritchie MD, Pendergrass SA. Integrating clinical laboratory measures and ICD-9 code diagnoses in phenome-wide association studies. Pac Symp Biocomput. 2016;21:168–179.

15. Wei WQ, Bastarache LA, Carroll RJ, Marlo JE, Osterman TJ, Gamazon ER, Cox NF, Roden DM, Denny JC. Evaluating phecodes, clinical classification software, and ICD-9-CM codes for phenome-wide association studies in the electronic health record. PLoS One. 2017:12:e0175508.

16. Carroll RJ, Bastarache L, Denny JC. R PheWAS: data analysis and plotting tools for phenome-wide association studies in the R environment. Bioinformatics. 2014;30:2375–6.

17. O’Seaghdha CM, Wu H, Yang Q, Kapur K, Guessous I, Zuber AM, Kottgen A, Stoudmann C, Teumer A, Kutalik Z, Mangino M, Dehghan A, Zhang W, Eiriksdottir G, Li G, Tanaka T, Portas L, Lopez LM, Hayward C, Lohman K, Matsuda K, Padmanaghan S, Firsov D, Sorice R, Ulivi S, Brockhaus AC, Kleber ME, Mahajan A, Ernst FD, Gudnason V, Launer LJ, Mace A, Boerwinckle E, Arking DE, Tanikawa C, Nakamura Y, Brown MJ, Gaspoz JM, Theler JM, Siscovick DS, Psaty BM, Bergmann S, Vollenweider P, Vitart V, Wright AF, Zemunik T, Boban M, Kolcic I, Navarro P, Brown EM, Estrada K, Ding J, Harris Tb, Bandinelli S, Hernandez D, Singleton AB, Girotto G, Ruggiero D, d’Adamo AP, Robino A, Meitinger T, Meisinger C, Davies G, Starr JM, Chambers JC, Boehm BO, Winkelmann BR, Huang J, Murgia F, Wild SH, Campbell H, Morris AP, Franco OH, Hofman A, Uitterlinden AG, Rivadeneira F, Völker U, Hannemann A, Biffar R, Hoffmann W, Shin SY, Lescuyer P, Henry H, Schurmann C, SUNLIGHT Consortium; GEFOS Consortium, Munroe PB, Gasparini P, Pirastu N, Ciullo M, Geiger C, März W, Lind L, Spector TD, Smith AV, Rudan I, Wilson JF, Polasek O, Deary IJ, Pirastu M, Ferrucci L, Liu Y, Kestenbaum B, Kooner JS, Witteman JC, Nauck M, Kao WH, Wallaschofski H, Bonny O, Fox CS, Bochud M. Meta-analysis of genome-wide association studies identifies six new loci for serum calcium concentrations. PLoS Genet. 2013;9(9):e1003796.

18. Dewey FE, Gusarova V, Dunbar RL, O’Dushlaine C, Schurmann C, Gottesman O, McCarthy S, Van Hout CV, Bruse S, Dansky HM, Leader JB, Murray MF, Ritchie MD, Kirchner HL, Habegger L, Lopez A, Penn J, Zhao A, Shao W, Stahl N, Murphy AJ, Hamon S, Bouzelmat A, Zhang R, Shumel B, Pordy R, Gipe D, Herman GA, Sheu WHH, Lee IT, Liang KW, Guo X, Rotter JI, Chen YI, Kraus WE, Shah SH, Damrauer S, Small A, Rader DJ, Wulff AB, Nordestgaard BG, Tybjaerg-Hansen A, van den Hoek AM, Princen HMG, Ledbetter DH, Carey DJ, Overton JD, Reid JG, Sasiela WJ, Banerjee P, Shuldiner AR, Borecki IB, Teslovich TM, Yancopoulos GD, Mellis SJ, Gromada J, Baras A. Genetic and pharmacologic inactivation of ANGPTL3 and cardiovascular disease. N Engl J Med. 2017;377:211–221.

19. Squires PE, Jones PM, Younis MY, Hills CE. The calcium-sensing receptor and β-cell function. Vitam Horm. 2014;95:249–267.

20. Babinsky VN, Hannan FM, Ramracheya RD, Zhang Q, Nesbit MA, Hugill A, Bentley L, Hough A, Joynson E, Stewart M, Aggarwal A, Prinz-Wohlgenannt M, Gorvin CM, Kallay E, Wells S, Cox RD, Richards D, Rorsman P, Thakker RJ. Mutant mice with calcium-sensing receptor activation have hyperglycemia that is rectified by calcilytic therapy. Endocrinol. 2017;158:2486–2502.

21. Graca JA, Schepelmann M, Brennan SC, Reens J, Chang W, Yan P, Toka H, Riccardi D, Price SA. Comparative expression of the extracellular calcium-sensing receptor in the mouse, rat and human kidney. Am J Physiol Renal Physiol. 2016;310:F518–F533.

22. Reid IR, Birstow SM, Bolland MJ. Calcium and cardiovascular disease. Endocrinol Metab (Seoul). 2017;32:339–349.

23. Smajilovic S, Tfelt-Hansen J. Calcium acts as a first messenger through the calcium-sensing receptor in the cardiovascular system. Cardiovasc Res. 2007;75:457–467.

24. Alam MU, Kirton JP, Wilkinson FL, Towers E, Sinha S, Rouhi M, Vizard TN, Sage AP, Martin D, Ward DT, Alexander MY, Riccardi D, Canfield AE. Calcification is associated with loss of functional calcium-sensing receptor in vascular smooth muscle cells. Cardiovasc Res. 2009;81:260–268.

25. Oh J, Beckmann J, Bloch J, Hettgen V, Mueller J, Li L, Hoemme M, Gross ML, Penzel R, Mundel P, Schaefer F, Schmitt CP. Stimulation of the calcium-sensing receptor stabilizes the podocyte cytoskeleton, improves cell survival, and reduces toxin-induced glomerulosclerosis. Kidney Int. 2011;80:483–492.

26. Babinsky VN, Hannan, FM, Youhanna SC, Maréchal C, Jadoul M, Devuyst O, Thakker RV. Association studies of calcium-sensing receptor (CaSR) polymorphisms with serum concentrations of glucose and phosphate, and vascular calcification in renal transplant recipients. PLoS One. 2015;10:30119459.

