## Supplemental Information for "Calcium Sensing Receptor Common Variants Influence the Effects of Serum Calcium on Coronary Artery Disease Risks"

***Supplemental Figures***


Figure S1. ExWAS regional association plot for SNP rs1801725 Ala986Ser), using total serum $Ca^{2+}$ measures for 51,289 individuals…..……………………………………………………..4
Figure S2. *CASR* genotype dependence of CAD comorbidities……….………………………….5
Figure S3. Lipid quantitative measures by *CASR* genotype…..…………………………….………...7


***Supplemental Tables***


Table S1. Demographics and Clinical Characteristics of Individuals in the DiscovEHR cohort……………………………………………………………………………………………………8
Table S2. Statistical analysis of tri-locus haplotype associations with serum calcium………...9
Table S3. Statistical analysis of SNPs and age, sex and BMI with CAD risk as a function of serum calcium.……………………………………………………………………………………….10
Table S4. Statistical analysis of age, sex and BMI with CAD risk as a function of serum calcium for CaSR haplotypes......………………………………………………………………..11

***References***……………………………………………………………………………..…….12

### *Investigators*

Diane T. Smelser, PhD,[1]§, Fadil M. Hannan, DPhil,[2,3]§, Raghu P.R. Metpally, PhD,[1], Sarathbabu Krishnamurthy, MS,[1], David J. Carey, PhD,[1], Rajesh V. Thakker, MD,[3], Gerda E. Breitwieser, PhD,[1], on behalf of Regeneron Genetics Center,[4].

[1]Weis Center for Research, Geisinger, Danville, PA, USA

[2]Nuffield Department of Women's & Reproductive Health, University of Oxford, Oxford, UK

[3]Academic Endocrine Unit, Radcliffe Department of Medicine, University of Oxford, Oxford, UK

[4]Regeneron Genetics Center, Tarrytown, NY, USA

§DTS and FMH contributed equally.

**[4]Regeneron Genetics Center Banner Author List (in alphabetical order)**

Goncalo Abecasis, Ph.D., Xiaodong Bai, Ph.D., Suganthi Balasubramanian, Ph.D., Nilanjana Banerjee, Ph.D., Aris Baras, M.D., Christina Beechert, Andrew Blumenfeld, Michael Cantor, M.D., Giovanni Coppola, M.D., Yating Chai, Ph.D., Amy Damask, Ph.D., Colm O'Dushlaine, Ph.D., Aris Economides, Ph.D., Gisu Eom, Caitlin Forsythe, M.S., Jan Freudenberg, M.D., Erin D. Fuller, Claudia Gonzaga-Jauregui, Ph.D., Nehal Gosalia, Ph.D., Zhenhua Gu, M.S., Lauren Gurski, Paloma M. Guzzardo, Ph.D., Lukas Habegger, Ph.D., Young Hahn, Alicia Hawes, B.S., Julie Horowitz, Ph.D., Marcus B. Jones, Ph.D., Shareef Khalid, Michael Lattari, Alexander Li, Ph.D., Nan Lin, Ph.D., Daren Liu, Alexander Lopez, M.S., Kia Manoochehri, Jonathan Marchini Ph.D., Anthony Marcketta, Evan K. Maxwell, Ph.D., Shane McCarthy, Ph.D., Lyndon J. Mitnaul, Ph.D., John D. Overton, Ph.D., Charles Paulding, Ph.D., John Penn, Kavita Praveen, Ph.D., M.S., Jeffrey G. Reid, Ph.D., Thomas D. Schleicher, M.S., Claudia Schurmann, Ph.D., Maria Sotiropoulos Padilla, M.S., Karina Toledo, Louis Widom, Sarah E. Wolf, M.S., Manasi

Pradhan, M.S., Alan Shuldiner, M.D., Jeffrey C. Staples, Ph.D., Dylan Sun, Tanya Teslovich, Ph.D., Ricardo H. Ulloa, Cristopher Van Hout, Ph.D., Ashish Yadav, M.S., Bin Ye.

### *Supplemental Figures*

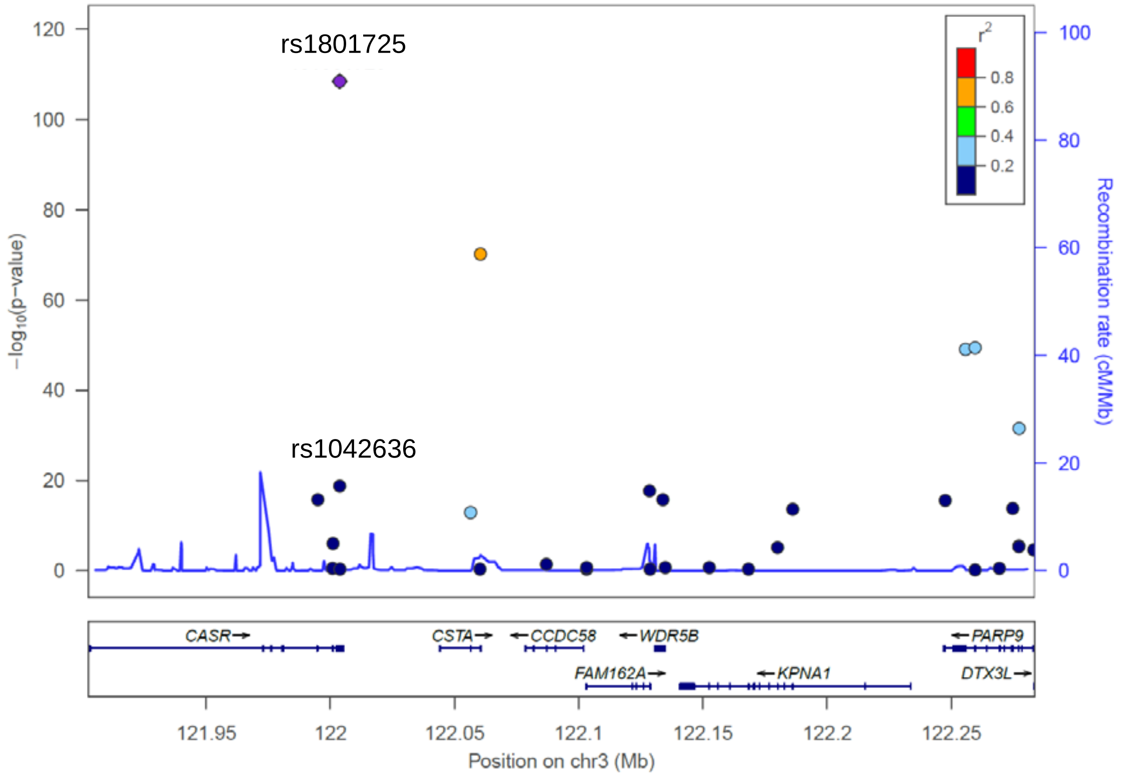


**Figure S1. ExWAS regional association plot for SNP rs1801725, using total serum calcium measures for 51,289 individuals.** For the single-marker Exome-Wide Association Study (ExWAS), all bi-allelic variants from exome sequencing data with missingness rates less than 2%, Hardy-Weinberg equilibrium p-values greater than 1.0E-07, and minor allele frequency (MAF) greater than 1% were analyzed. Samples were also filtered by applying a call rate of 98%. We used mixed linear model-based association analysis (GCTA-MLMA)[1] to test for associations between single variants and serum calcium levels, fitting a genetic relatedness matrix (GRM; constructed from markers in approximate linkage equilibrium with MAF greater than 0.1%) as a random-effects covariate. MLMA analysis will account for relatedness and population structure from ancestry.[2] Plot generated using stand-alone locus zoom tool.[3]

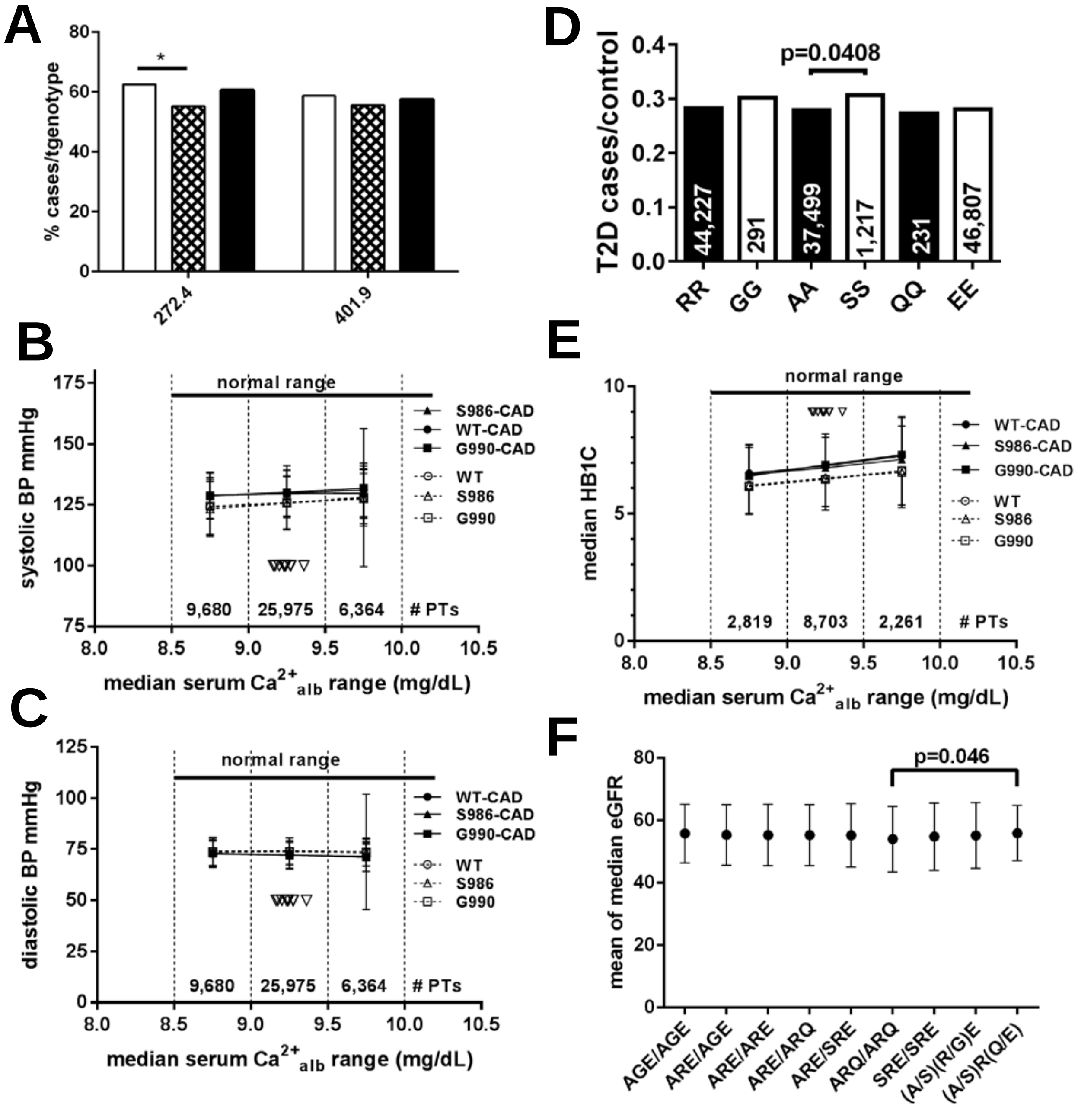


**Figure S2.** ***CASR*** **genotype dependence of CAD comorbidities. A**. Individuals homozygous for S986 (open bar), G990 (crosshatched bar) or E1011 (black bar) were examined scored for ≥ 3 call for dyslipidemia (ICD9 272.4) or essential hypertension (ICD9 401.9). There was a marginally significant difference between cases between S986 and G990 (Chi-squared with Yates' test, *p<0.05). **B.-C.** Individuals having measures in the EHR for blood pressure, were grouped as a function of $LM_{EHR}$ median $Ca^{2+}_{alb}$, and plotted by genotype for those with CAD or without (controls). There were no significant differences by genotype or serum $Ca^{2+}$. Inverted arrowheads mark mean serum $Ca^{2+}$ levels for haplotypes listed in Table 1. **D.** *CASR* SNP contributions to type 2 diabetes (T2D). Cases are grouped by SNP, with only homozygous individuals plotted (other positions in the haplotype were reference). Numbers of patients with each genotype are indicated; significance was determined by Fisher's exact test. **E.** Hemoglobin A1c median values for *CASR* genotypes of CAD and non-CAD patients are plotted. There were no significant differences by genotype. **F.** Means of median eGFRs were plotted against *CASR* haplotypes. Significant differences among haplotypes are noted.

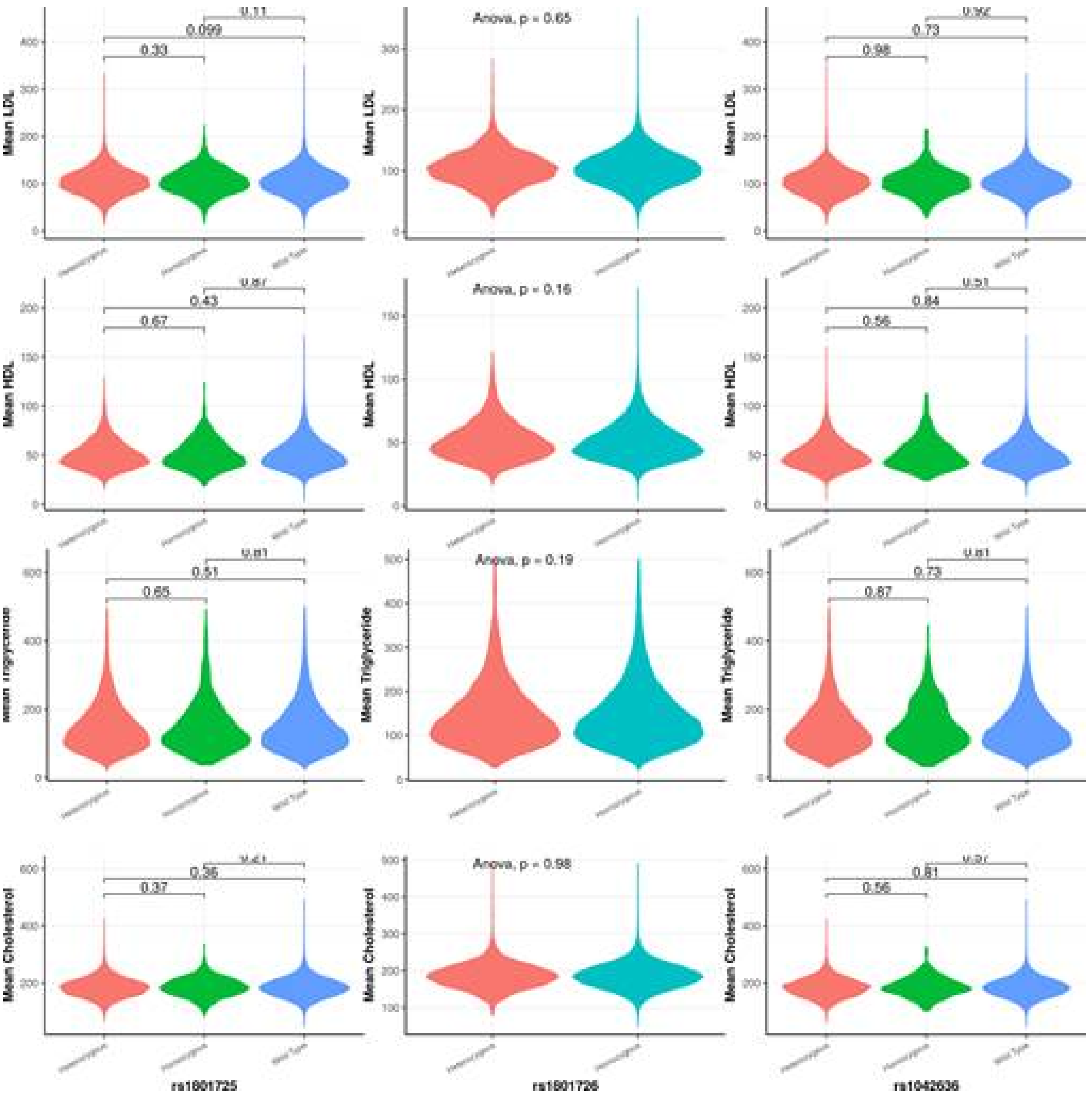


**Figure S3. Lipid quantitative measures by *CASR* haplotype.** Lipid levels plotted by haplotype for patients having measures in their EHR; there were no significant differences among haplotypes.

### *Supplemental Tables*

**Table S1. Demographics and Clinical Characteristics of Individuals in the DiscovEHR cohort.**

| **Basic Demographics** | **DiscovEHR Sequenced Patients[1]** |
|---|---|
| N | 51,289 |
| # of individuals <18 yrs of age | 963 |
| Female, N (%) | 30,290 (59) |
| Median Age, yr | 61 (48-72) |
| Median BMI, $kg/m^2$ | 30(26-36) |
| Median Years of EHR data | 14 (11-17) |
| Median medication orders/patient | 129 (37-221) |
| Median lab results/patient | 658 (197-1,119) |
| # PTs with median $Ca^{2+}$-albumin, N, (%) | 42,466 (83%) |

[1]Values are expressed as median (interquartile range).

[2]Abbreviations, EHR, electronic health record, BMI, GHS, Geisinger Health System, BMI, body mass index.

**Table S2. Statistical analysis of tri-locus haplotype associations with median $LM_{EHR}$ serum $Ca^{2+}_{alb}$.** Defined haplotypes were compared to all other haplotypes, numbers of patients in each group indicated in parentheses in first column. Associations between the haplotypes and the continuous variables was performed with an analysis of variance run using general linear models (GLM), controlling for age and sex. The matrix of P values for pairwise comparisons of each haplotype was performed with least square means. All statistical analyses were performed with SAS 9.4 for Windows (SAS Institute Inc, Cary, NC).

| | **ARE/ARE** | **ARE/ARQ** | **ARQ/ARQ** | **ARE/AGE** | **AGE/AGE** | **ARE/SRE** | **SRE/SRE** |
|---|---|---|---|---|---|---|---|
| **ARE/ARE (16,889)** | X | **0.0017** | 0.8383 | **<0.0001** | 0.7371 | 0.3033 | **<0.0001** |
| **ARE/ARQ (1,973)** | **0.0017** | X | 0.4529 | **<0.0001** | 0.1969 | 0.06 | **<0.0001** |
| **ARQ/ARQ (54)** | 0.8383 | 0.4529 | X | 0.6659 | 0.987 | 0.7177 | 0.0976 |
| **ARE/AGE (3,296)** | **<0.0001** | **<0.0001** | 0.6659 | X | 0.422 | 0.9839 | **<0.0001** |
| **AGE/AGE (148)** | 0.7371 | 0.1969 | 0.987 | 0.422 | X | 0.5898 | **0.0032** |
| **ARE/SRE (6,702)** | 0.3033 | 0.06 | 0.7177 | 0.9839 | 0.5898 | X | **0.0007** |
| **SRE/SRE (721)** | **<0.0001** | **<0.0001** | 0.0976 | **<0.0001** | 0.0032 | 0.0007 | X |

**Table S3. Statistical analyses of SNP, age, sex, BMI associations with CAD risk as a function of median $LM_{EHR}$ serum $Ca^{2+}_{alb}$.** Calcium values were categorized into three groups for comparison: (A) 8.5-9.0 mg/dL, (B) 9.0-9.5 mg/dL and (C) 9.5-10.0 mg/dL. These groups were compared in three separate analyses: A vs. B, A vs. C and B vs. C, using logistic regression for age, sex (male vs female), and BMI (age/BMI at date when median serum $Ca^{2+}_{alb}$ data was obtained). Odds ratio estimates and 95% Wald confidence limits are given, along with the effect size, $\beta$, and p-values. Analysis was performed with SAS 9.4 (SAS Institute, Cary, NC). Statistical significance is indicated, *p<0.05, **p<0.01; ***p<0.001, ****p<0.0001. Bold denotes significant associations.

| ***CASR* SNP** | **Risk factor** | **A vs B** | **A vs C** | **B vs C** |
|---|---|---|---|---|
| **Glu1011Gln** | **SNP** | 1.86 (0.471-7.732) | 0.255 (0.05-1.293) | **0.157 (0.038-0.65)*** |
| | **age** | 1.114 (1.043-1.19)** | 1.102 (10.25-1.185)** | 1.105 (1.033-1.182)** |
| | **sex** | 149 (0.37-5.92) | 4.26 (0.82-22.2) | 3.86 (0.99-15.1) |
| | **BMI** | 1.05 (0.96-1.15) | 1.08 (0.98-1.22) | 1.114 (1.014-1.23)* |
| **Ala986Ser** | **SNP** | **0.878 (0.798-0.966)**** | **0.726 (0.633-0.833)****** | **0.862 (0.769-0.967)*** |
| | **age** | 1.059 (10.56-1.063)**** | 1.059 (1.054-1.064)** | 10.57 (1.054-1.061)**** |
| | **sex** | 3.16 (2.9-3.43)**** | 3.29 (2.9-3.73)**** | 3.06 (2.82-3.32)**** |
| | **BMI** | 1.03 (1.01-1.023)**** | 1.03 (1.02-1.04)**** | 1.01 (1.01-1.02)**** |
| **Arg990Gly** | **SNP** | **0.912 (0.833-0.998)*** | **0.83 (0.733-0.939)**** | **0.905 (0.819-1.0)*** |
| | **age** | 1.06 (1.055-1.061)**** | 1.057 (1.051-1.062)**** | 1.055 (1.051-1.059)**** |
| | **sex** | 3.2 (2.9-3.5)**** | 3.7 (3.22-4.26)**** | 3.08 (2.82-3.37)**** |
| | **BMI** | 1.02 (1.01-1.02)**** | 1.03 (1.02-1.04)**** | 1.01 (1.005-1.059)**** |

**Table S4. Statistical analysis of age, sex, BMI associations with CAD risk as a function of median $LM_{EHR}$ serum $Ca^{2+}_{alb}$ for *CASR* haplotypes.** Each SNP in the haplotype was tested separately. Three ranges of median albumin-adjusted serum $Ca^{2+}$ values were generated for comparison: (A) 8.5-9.0 mg/dL, (B) 9.0-9.5 mg/dL, and (C) 9.5-10.0 mg/dL. These groups were compared in three separate analyses: A vs. B, A vs. C, and B vs. C, using logistic regression for age, sex (male vs female), and BMI (age/BMI at date when median serum $Ca^{2+}_{alb}$ data was obtained). Odds ratio estimates and 95% Wald confidence limits are given, and p-values are indicated as *p<0.01 or **P<0.0001. Analysis was performed with SAS 9.4 (SAS Institute, Cary, NC). (N) indicates number of individuals with tested haplotype in the designated median $LM_{EHR}$ serum $Ca^{2+}_{alb}$ ranges.

| ***CASR* Haplotype** | **Risk factor** | **A vs B** | **A vs C** | **B vs C** |
|---|---|---|---|---|
| **Ala986 Arg990 Glu1011** | **(N)** | **21,708** | **9,821** | **19,201** |
| | **age** | 1.059 (1.055-1.063)** | 1.057 (1.051-1.063)** | 1.056 (1.052-1.060)** |
| | **sex** | 3.09 (2.0-3.41)** | 3.55 (3.051-4.137)** | 2.97 (2.69-3.28)** |
| | **BMI** | 1.016 (1.009-1.023)** | 1.028 (1.018-1.037)** | 1.01 (1.004-1.017)* |
| **Ala/Ser+Ser/Ser986 Arg990 Glu1011** | **(N)** | **Alt=8,141; Ref=21,708** | **Alt=3,839; Ref 9,821** | **Alt=8,428; Ref=19,201** |
| | **age** | 1.059 (1.056-1.063)** | 1.058 (1.053-1.063)** | 1.057 (1.054-1.061)** |
| | **sex** | 3.161 (2.91-3.44)** | 3.292 (2.901-3.737)** | 3.061 (2.82-3.322)** |
| | **BMI** | 1.06 (1.011-1.022)** | 1.028 (1.021-1.037)** | 1.014 (1.008-1.019)** |
| **Ala986 Arg/Gly+Gly/Gly990 Glu1011** | **(N)** | **Alt-4,261; Ref=21,708** | **Alt=2,010; Ref=9,821** | **Alt=3,451; Ref=19,201** |
| | **age** | 1.057 (1.053-1.06)** | 1.057 (1.051-1.062)** | 1.055 (1.051-1.059)** |
| | **sex** | 3.21 (2.93-3.52)** | 3.714 (3.224-4.278)** | 3.082 (2.82-3.38)** |
| | **BMI** | 1.015 (1.009-1.021)** | 1.028 (1.019-1.037)** | 1.011 (1.005-1.017)* |
| **Ala986 Arg990 Glu/Gln+Gln/Gln1011** | **(N)** | **Alt=123; Ref=21,708** | **Alt=67; Ref=9,821** | **Alt=112; Ref=19,201** |
| | **age** | 1.059 (1.055-1.063)** | 1.057 (1.052-1.063)** | 1.056 (1.052-1.06)** |
| | **sex** | 3.079 (2.79-3.396)** | 3.55 (3.052-4.132)** | 2.97 (2.69-3.27)** |
| | **BMI** | 1.016 (1.009-1.023)** | 1.028 (1.019-1.038)** | 1.011 (1.004-1.018)* |

### REFERENCES


1. Yang J, Lee SH, Goddard ME, Visscher PM. GCTA: a tool for genome-wide complex trait analysis. *Am J Hum Genet.* 2011;88:76-82.
2. Yang J, Zaitlen NA, Goddard ME, Visscher PM, Price AL. Advantages and pitfalls in the application of mixed-model association methods. *Nat Genet.* 2014;46:100-106.
3. Pruim RJ, Welch RP, Sanna S et al. LocusZoom: regional visualization of genome-wide association scan results. *Bioinformatics.* 2010;26:2336-2337.
